# The preparation of cyclopentadecanone and cyclopentadecanolide from *Malania oleifera Chum* oil

**DOI:** 10.1101/622423

**Authors:** Pin Liu, Xiongmin Liu, Fang Lai, Li Ma, Weiguang Li, Pinxian Huang

**Affiliations:** (College of Chemistry and Chemical Engineering, Guangxi University, Nanning, China); Department of Science and Technology, Guangxi University for Nationalities, Nanning 530006, China; Nanning Pinliu Biotechnology Co., Ltd, Nanning 530028, China

**Keywords:** *Malania oleifera Chum* oil, cyclopentadecanone, cyclopentadecanolide, preparation

## Abstract

The macrocyclic musks are of central importance in the fragrance industry. The preparation of cyclopentadecanone and cyclopentadecanolide from *Malania oleifera Chum* oil were investigated. Preparation method of cyclopentadecanone is three steps process: first fatty acid double bonds of *Malania oleifera Chum* oil are cut off by ozonolysis; second ozonides are converted into methyl esters by esterification; afterwards cyclopentadecanone was achieved from through acyloin condensation, reduction and separation with 38.5 % overall yield. The preparation coyfclopentadecanolide is two step process: first by ozonolysis triglycerides of ω-hydroxycarboxylic acids synthesized from malania oleifera chum oil by ozonolysis; and it were catalytically transformed to macrocyctic lactones, and a 63% yield of cyclopentadecanolide was obtained. The effect of different ester groups on cyclization of α,ω-difatty acid alkyl ester, and effect of catalysts on cyclization of ω-hydroxycarboxylic acid triglyceride were investigated.

## 1. Introduction

Cyclopentadecanolide and cyclopentadecanone are two important macrocyclic musk[1], and widely used in perfumes, cosmetics, food, and medical. Cyclopentadecanolide exists in many plants [2, 3], but its content is very low, this is difficult to extract and separate cyclopentadecanolide. Several preparation methods of cyclopentadecanolide have been developed [4, 5, 6, 7]. Ookoshi [8] has been reported that the macrolactonization of x-hydroxyalkanoic acid in a highly concentrated solution is catalyzed by dealuminated HY zeolite. The HZSM-5 zeolite is one of the solid acids which have been widely used as viable alternatives to conventional acids in esterification reactions [9, 10, 11]. Lai [12] has been reported that macrolactonization of methyl 15-hydroxypentadecanoate to cyclopentadecanolide over Mo-Fe/HZSM-5 catalyst. The important intermediate of synthetic cyclopentadecanolide is 15-hydroxyalkanoic acid. However, the manufacture of 15-hydroxyalkanoic acid is complex and difficult when synthesized by chemical methods. Cyclopentadecanone occurs, along with cyclopentadecanol in the secretion of the North American musk rat. McGinty reported the fragrance material review on cyclopentadecanone [13]. Many synthetic routes of cyclopentadecanone have been reported [14, 15, 16, 17, 18, 19]. The synthesis of 1,15-pentadecanedioate is complex, if chemical synthesis is used from petroleum products.

*Malania oleifera Chum et S. Lee* (simple name: *Malania oleifera Chum*) is a wild woody plant, mainly distributed in Guangxi and Yunnan Province of China [20]. Its fruits contains oils and fats [21] (50-55% by weight) in which the main component is 15-tetracosenoic acid (nervonic acid) (40-50% by weight). The aforementioned compound is a good candidate for synthesizing macrocyclic musks, such as cyclopentadecanolide [22, 23]. However, it is difficult to separate and purify 15-tetracosenoic acid from the fatty acids mixture of *Malania oleifera Chum* fat [24], The yield of 15-tetracosenoic acid is only 15%. Therefore, the cost of cyclopentadecanolide is relatively high when 15-tetracosenoic acid is used as raw material.

In the present work, the purpose is to make efficient use of natural plant resources, the preparation of cyclopentadecanolide and cyclopentadecanone were explored for improve the efficiency of 15-tetracosenoic acid and reduce the cost. It is helpful for the commercial use of *Malania oleifera Chum*. This is very interesting method for the way to cut off fat double bond directly and then macrocyclization. Preparation of cyclopentadecanone and cyclopentadecanolide from *Malania oleifera Chum* oil is very meaningful for preparation methods and technologies, because fatty acid glycerides directly synthesize macrocyclic flavors are rarely reported in the literature.

## 2. MATERIALS AND EXPERIMENTS

### 2.1. Materials and apparatus

*Malania oleifera Chum* oil was extracted from fruit of *Malania oleifera Chum et S. Lee* harvested in Guangxi province of China. Plant species identified by Professor Lai Jiaye, botany expert (College of Forestry, Guangxi University).

Standard cyclopentadecanone was purchased from Aldrich Co. (USA), purity: 98.0 %. Standard cyclopentadecanolide was purchased from Askrich Chemical Company Inc. (Japan), purity: 98%.

IR spectra were recorded by a SHIMAZHU FT-IR8400S spectrometer (Japan). Mass spectra were determined by SHIMAZHU GC-MS/QP5050A (Japan). The quantity of cyclopentadecanone was analyzed by SHIMAZHU GC - 16A (Japan). Bruker 600 MHz NMR.

### 2.2. Extraction of *Malania oleifera Chum* oil and preparation of fatty acid methyl esters

The extraction followed Soxhlet method using 50 g sample of *Malania oleifera Chum*. It included the crushed *Malania oleifera Chum* and 8 h Soxhlet reflux in petroleum ether of boiling range 60-90 °C. Preparation of fatty acid methyl esters used a slightly modified method based on Simoneau and Vicario et al [25, 26].

### 2.3. Analysis of fatty acid methyl esters

SHIMAZHU GC-MS/QP5050A (Japan) equipped with USA J&W Co. DB-1 column (30.0 m×0.25 mm×0.25 m) was used for compound identification. Helium was employed as carrier gas at a constant flow rate of 1.5mLmin^−1^. Initial oven temperature was set at 150 °C, held for 1min, ramped at 4°C min^−1^ to 270 °C and held for further 5min with heated capillary transfer line maintained at 270°C. Splitless injection was carried out at 270 °C and 0.2μL of sample was injected. In the GC/MS full scan mode, m/z 40 to m/z 450 was recorded. Chromatographic peaks were identified with NIST mass spectral data library and the retention times were compared with standard compounds listed in NIST 2008 Mass Spectral Libraries V2.2.

The consists of oils and fats 53.2 %(w/w), in which the fats mainly contain 15-tetracosenic acid 46.7 %, 9-octadecenoic acid 27.9 %, and erucic acid 12.5 %, other fatty acids 12.9 %.

### 2.4. Preparation of cyclopentadecanone

A solution of *Malania olceifera Chum* oil (40.0 g) in hexane (300 mL) and acetic acid (90 mL) was ozonized at 0 °C for 4 hr, and then H_2_O_2_ (30 %, 40 mL) was added dropwisely for another 3 hr period at room temperature. The reaction mixture was added to ice water, filtered and washed with water to obtain solid. The solid was dried to give 30 g mixed products P1.

P1 30 g, Methanol (270 g), and sulfuric acid (6 g) was mixed and refluxed for 4 hr. The mixture was cooled, extracted with diethyl ether (100 mL 2 times), distilled to provide 30g mixture of α,ω-difatty acid methyl ester P2.

P2 30g in xylene (40 mL.) was added to pulverized sodium (10g.) in refluxing xylene (500 mL) under nitrogen during 1 hr. The mixture was refluxed for 1 hr. Then, 150 mL ethanol was added slowly to the reactor at 80°C, after cooling to room temperature. Acetic acid(100 mL) was added to reactor, followed by 150 mL water. The xylene solution was separated from reaction mixture, and then treated with water. Xylene was evaporated under reduced pressure. The residue was distilled to give 16.0 g acyloin.

Hydrochloric acid (16 mL) was added to the mixture of crude acyloin (16.0 g.) and Zn powder (4g) during 1 hr at 110°C. After the reaction maintained for another 30 min, the mixture was cooled to room temperature, extracted with benzene (200 mL), and washed with water.

The mixture was distilled, separated by vacuum distillation (or vacuum distillation of glycerin as entrainer) and recrystalled with ethanol to obtain cyclopentadecanone 16.0g with purity of 97.4%.

### 2.5. Preparation of cyclopentadecanolide

A solution of Malania olceifera Chum oil (200.0 g) in hexane (200 mL) and ethanol (200 mL) was ozonized at 0 °C for 4 hr, and then reactant was added dropwisely 50 g potassium borohydride was dissolved in 500mL aqueous solution at 10 °C for 3 hr. Then neutralize to neutrality with hydrochloric acid for 2 hr, filteration, washing with water for 3 times, and drying to obtain ω-hydroxycarboxylic acid triglyceride 135 g.

Put ω-hydroxycarboxylic acid triglyceride (5) (100 g) and catalyst into 1000 mL three bottles, then slowly add glycerol and heating up to make glycerol distillate at vacuum degree is <755 mmHg, and mixture products 75 g was obtained when reaction is 40 hr. The mixture was fractionating by distillation and recrystallized by ethanol, and cyclopentadecanolide **4**7 g were obtained.

### 2.6. Analytical of cyclopentadecanone and cyclopentadecanolide

The structures of cyclopentadecanone and cyclopentadecanolide were determined by IR (KBr), GC-MS, ^1^H NMR (600 MHz, CDCl_3_) and ^13^C NMR (600 MHz, CDCl_3_), and compared with their standards.

## 3. RESULTS AND DISCUSSION

### 3.1. Preparation method of cyclopentadecanone

Although there are many preparation methods for cyclopentadecanone, the typical technological process is shown in the figure 1.

**Fig.1.**
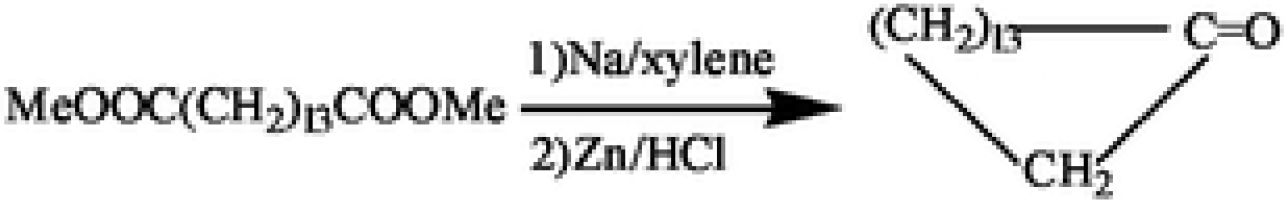
typical preparation method of cyclopcntadccanone.

The production costs of MeOOC(CH_2_)_13_COOMe or HOOC(CH_2_)_13_COOH are high, in the preparation of cyclopentadecanone. Interestingly, the content of 15-tetracosenoic acid is 46.7 % in *Malania oleifera Chum* oil, it is a good raw material for preparing cyclopentadecanone. In this paper, preparation of the cyclopentadecanone from *Malania oleifera Chum* oil was described and displayed in the figure 2.

**Fig.2.**
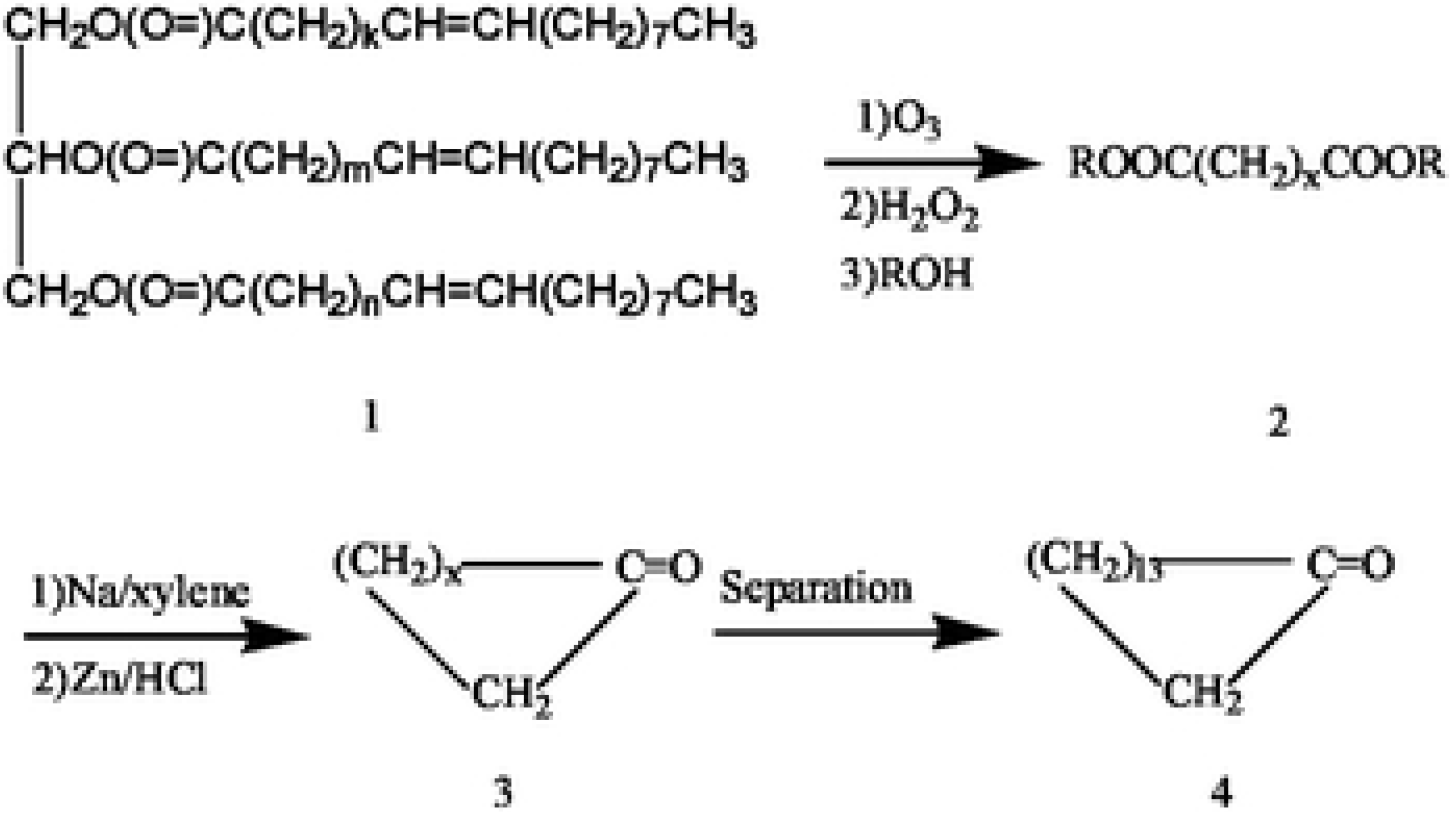
preparation method of cyclopentadecanonc from *Malania oleifera Chum* oil.

The intermediate methyl ester was produced from the starting material 1 *via* a three steps process which consists of ozonolysis oxidation and esterification. The mixed product 3 was achieved from 2 through acyloin condensation and reduction. Afterwards, mixed product 3 was separated to product cyclopentadecanone 4 by distillation its total yield (in terms of tetracosolenicacid-15) is 38.5 %.

Comparing the prepared cyclopentadecanone with the standard cyclopentadecanone, IR (KBr), GC-MS, ^1^H NMR (600 MHz, CDCl_3_) and ^13^C NMR (600 MHz, CDCl_3_) are identical.

### 3.2. The effect of groups on the yield of cyclopentadecanone

In figure 2, when group Me is replaced by other groups, does it affect the yield? The effect of groups on the yield of cyclopentadecanone was investigated and the result was shown in Table 1.

**Table 1.** the effect of groups on the yield.

| Ester groups | Yields* of cyclopentadecanone (%) |
| --- | --- |
| methyl | 32.7 |
| ethyl | 38.5 |
| propyl | 31.1 |
| butyl | 21.0 |

\*in terms of 15-tetracosenic acid

Table 1 shows that the yield of cyclopentadecanone from ethyl ester was the highest (38.5%) among these ester groups.

### 3.3 Preparation method of cyclopentadecanolide

The preparation method of cyclopentadecanolide from *Malania oleifera Chum* oil is shown in figure 3 by Guo [22].

**Fig.3.**
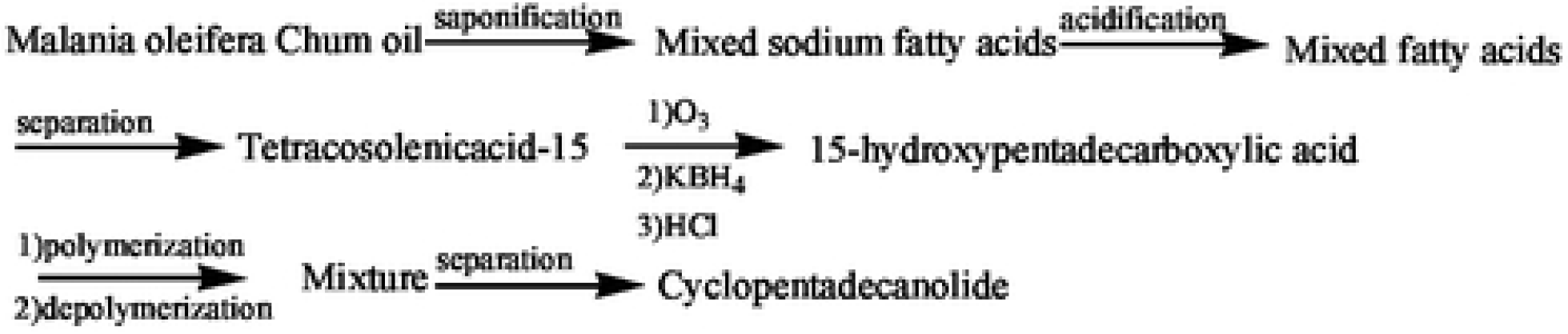
preparation method of cyclopcntadec anolide from *Malania oleifera Chum* oil.

Although on chemical synthesis, the preparation of mixed fats of is simple from *Malania oleifera Chum* oil, but separation of 15-tetracosenic acid is difficult to use crystallization method, because15-tetracosenic acid, 9-octadecenoic acid and erucic acid properties are similar in mixed fats. The yield of 15-tetracosenic acid (purity 95%) is only about 10% from *Malania oleifera Chum* oil. Therefore, the utilization of 15-tetracosenic acid is low. The simple preparation method of cyclopentadecanolide from *Malania oleifera Chum* oil was described in the figure 4.

**Fig.4.**
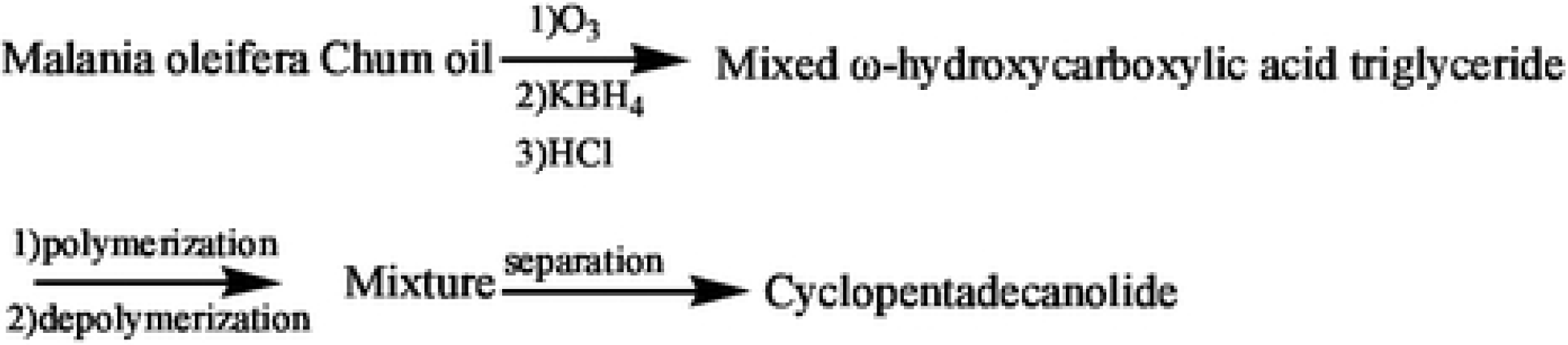
The simple preparation method of cyclopcntadec anolide from *Malania oleifera Chum* oil.

In figure 4, the intermediate mixed ω-hydroxycarboxylic acid triglyceride was produced from the starting material of *Malania oleifera Chum* oil via two process which consists of ozonization and reduction reaction. Afterwards, the desired product cyclopentadecanolide with 63 % overall yield was achieved from mixed ω-hydroxycarboxylic acid triglyceride through cyclization and separation. This is a very short technological route, and it is easy to industrialize.

Comparing the prepared cyclopentadecanolide with the standard cyclopentadecanolide, IR (KBr), GC-MS, ^1^H NMR (600 MHz, CDCl_3_) and ^13^C NMR (600 MHz, CDCl_3_) are identical.

### 3.4. Effect of catalyst on cyclopentadecanolide

The catalyst is important for the yield of cyclization reaction, effect of catalysts on cyclization of ω-hydroxycarboxylic acid triglyceride were investigated. The result was shown in table 2.

**Table 2.** Effect of catalysts on yields of cyclization.

| catalysts on | Yields of cyclopentadecanolide (%)* |
| --- | --- |
| NaOH | 31.0 |
| $\text{CH}_3\text{ONa}$ | 52.0 |
| $\text{CH}_3\text{ONa}/ \text{NaOH}$ | 63.0 |

\*in terms of 15-tetracosenic acid

Table 2 shows that the yield of cyclopentadecanolide is 63% when using mixed catalyst CH_3_ONa/ NaOH, it is high compared with catalyst single NaOH or CH_3_ONa. This synthetic route is short and has the value of industrial application.

## 4. CONCLUSIN

*Malania oleifera Chum* oil of plant resources are used to prepare musk perfume. The preparation method of cyclopentadecanone and cyclopentadecanolide were developed. Preparation method of cyclopentadecanone is three steps process which consists of ozonization, oxidation and esterification, and total yield is 38.5 % when in terms of 15-tetracosenic acid. The effects of ester groups on cyclopentadecanone were explored. This is a technological route of industrial value. Preparation method of cyclopentadecanolide is three steps process which consists of ozonization and reduction reaction, cyclization, separation and 63% yield of cyclopentadecanolide was obtained. The effect of catalysts on cyclization of ω-hydroxycarboxylic acid triglyceride was investigated. It is a very short technological route, and it is easy to industrialize.

## ACKNOWLEDGEMENTS

This work was supported by National Natural Science Foundation of China (21776050), Major science and technology special project in Guangxi (AA17204065-20), Science Foundation of Guangxi Education Department of China (2019KY0179), and Science Foundation of Guangxi University for Nationalities of China (2017MDQN004, XTCX201706).

